# Hypomethylation may drive *CLPB* overexpression which predicts poor survival outcomes in breast cancer

**DOI:** 10.1101/2025.10.12.681854

**Authors:** Sena Sonmez, Senem Noyan, Tolga Acun

## Abstract

**Background:** CLPB has oncogenic properties in leukemia and melanoma. We aimed to reveal the subtype-specific expression of the *CLPB* gene in breast cancer, its effect on survival, and genetic/epigenetic alterations that might regulate the expression.

**Methods:** Expression levels and prognostic signatures of *CLPB* gene in breast cancer patients were evaluated by using in-silico tools. RT-PCR and Q-RT-PCR methods were used to examine *CLPB* expression in cell lines. Genetic alterations of *CLPB* were revealed by employing Sanger Cosmic database. Combined bisulfite restriction analysis and UALCAN tool were used to examine the methylation status of the *CLPB* promoter in cell lines and samples, respectively.

**Results:** *CLPB* expression is significantly higher in all breast cancer subtypes compared to normals, and it is associated with worse survival values. Genetic alterations of *CLPB* are rare. Cell lines express *CLPB* gene at moderate to high levels. The promoter region of *CLPB* is hypomethylated in samples and cell lines.

**Conclusions:** We suggest *CLPB* gene as a therapeutic target and biomarker candidate in breast cancer. The rarity of genetic alterations suggests that they do not affect *CLPB* regulation, but promoter hypomethylation would be one of the reasons for its high expression in breast cancer.

## Introduction

Despite many advanced strategies developed for prevention, diagnosis, and treatment, breast cancer is still the most common type of cancer among women worldwide and can be challenging to treat [1]. Breast cancer can develop in the milk ducts or the milk-producing lobules, named as ductal and lobular carcinomas, respectively. Invasive ductal carcinoma is the most common type (70-80%) among all breast cancer cases. Breast cancer is very heterogeneous disease and typically classified into four subtypes based on the expression levels of estrogen receptor (ER), progesterone receptor (PR), and human epidermal growth factor receptor 2 (HER2): Luminal A (ER+, PR+, HER2–), luminal B (ER+, PR+, HER2+), HER2-enriched/positive (ER–, PR–, HER2+), and triple-negative (TNBC, basal-like) (ER–, PR–, HER2–) [2].

Currently, breast cancer treatment strategies are planned according to these subtypes. Patients with luminal subtypes are usually treated with endocrine therapy and/or chemotherapy. HER2-positive patients are treated with monoclonal antibodies and tyrosine kinase inhibitors. Patients with the basal-like subtype are more difficult to treat than those with other subtypes and typically receive chemotherapy [2].

Patients with luminal A subtype have the best prognosis with a lower incidence of cancer recurrence and higher survival rates among other subtypes. The prognosis of Luminal B is worse than Luminal A. The survival rates of the HER2-positive type have improved with the help of HER2-targeted therapies. However, they are more aggressive, grow faster, and have a worse prognosis compared to luminal tumors [3]. Patients with TNBC are more prone to poor prognosis and metastasis compared to patients with other subtypes. Cytotoxic chemotherapy drugs are frequently used for the treatment of TNBC [4].

So far, biomarkers and therapeutic targets have been identified that assist clinicians in breast cancer diagnosis, risk stratification, disease subtyping, selection of the most appropriate treatment, and survival analysis [5]. However, the identification of new biomarkers and therapeutic targets is important for the development of further diagnostic, prognostic, and treatment strategies for breast cancer subtypes.

CLPB (caseinolytic mitochondrial matrix peptidase chaperone subunit B) gene is located in chromosomal region 11q13.4 and encodes a mitochondrial chaperone protein. Proteomic analyses have shown that CLPB is located in the mitochondrial intermembrane space (IMS) of mitochondria, where it participates in ATP-dependent protein-protein interactions such as mitochondrial protein transport and quality control [6]. CLPB protein is important in maintaining mitochondrial cristae structure and regulating important mitochondrial biological processes. CLPB protein is overexpressed in acute myeloid leukemia (AML) cases, protects AML cells from apoptosis, and causes chemoresistance. A reduction in CLPB expression levels made AML cells more prone to apoptosis and more sensitive to the drug (venetoclax) by affecting the cristae structure of mitochondria and increasing mitochondrial oxidative stress. CLPB deprivation is thought to induce ATF4-driven mitochondrial stress response and increase pro-apoptotic signals. Reduced CLPB expression also results in impaired glycolysis, which suppresses cell proliferation. Leukemia development was significantly reduced and survival rates increased in mice that received “CLPB-deficient AML cells” intravenously compared to the control group [7]. Another study also shows that high CLPB expression is associated with poor prognosis in skin cutaneous melanoma (SCKM) and suppression of CLPB expression inhibited proliferation, invasion, and migration of melanoma cells [8]. High *CLPB* expression has also been reported to negatively affect survival in prostate cancer [9].

This background information suggests that overexpression of *CLPB* could be important in carcinogenesis and drug resistance, not only in leukemia and melanoma but also in other types of cancer. When we examined TCGA RNA-seq data with the UALCAN in-silico tool [10, 11], we found that *CLPB* expression is significantly high in many cancer types compared to normal tissues. These cancers include stomach, colon, head and neck, esophageal, liver, and lung cancers **(Supplementary Information Fig. 1)**.

In this study, we present comprehensive expression and survival statistics of *CLPB* gene in breast cancer by employing UALCAN [10, 11] and bc-GenExMiner [12, 13] in-silico tools. We have also documented the mutation and promoter methylation profile of *CLPB* in breast cancer samples and cell lines, which may affect the expression and/or function of *CLPB*.

## Material and methods

### Cell lines

BT-20 (triple negative A), SK-BR-3 (HER2+), MDA-MB-231 (triple negative B), ZR-75-1 (luminal A, ER+, PR+/-), MDA-MB-468 (triple negative A), BT-474 (luminal B, ER+, PR+, HER2+), MCF7 (luminal A, ER+, PR+) breast cancer cell lines [14] and MCF 10A (fibrocystic mammary epitel) cell line were cultured by following ATCC (American Type Culture Collection) guidelines.

### Multiplex Semi-quantitative RT-PCR

Total RNAs were extracted from breast cell lines by using the RNeasy Mini Kit (cat. no.: 74104; Qiagen, Germany). cDNAs were synthesized according to the Verso cDNA Synthesis Kit protocol (cat. no.: AB1453A; Thermo Fisher Scientific Inc., MA, USA). Multiplex semi-quantitative RT-PCR method was performed by using the *CLPB* and *TBP* (normalizer) [15] specific primers in the same reaction for 31 cycles. *CLPB* expressions of the cell lines were normalized in the second gel according to the band intensities of their *TBP* gene expressions in the first gel. The information about primers was given in **Supplementary Information Table 1**.

### Q-RT-PCR

Relative amounts of *CLPB* transcripts were determined by using iTaq™ Universal SYBR Green Supermix (cat. no.: 1725120) at the CFX96 Touch™ Real-Time PCR System (Bio-Rad Laboratories, Inc., CA, USA). *TBP* (TATA-box binding protein) gene was used as a normalizer [15]. The information about primers was given in **Supplementary Information Table 1**. Relative levels of transcripts were measured by Livak (2^-ddCT^) method [16].

### In-silico tools and databases

UALCAN data analysis portal could perform gene expression analysis based on RNA-seq data obtained from The Cancer Genome Atlas (TCGA). TPM (transcripts per million) values were used to generate boxplot charts. TPM values were also used to calculate the significance of gene expression differences between groups or subtypes with the t-test [10] (https://ualcan.path.uab.edu/). UALCAN could also perform promoter methylation analysis by using Infinium HumanMethylation450 BeadChip DNA methylation array data [17]. The average beta values of the samples, ranging from unmethylated (0) to fully methylated (1), were displayed on boxplot charts, and the statistical significance was calculated by t-test [11].

Bc-GenExMiner is a statistical mining tool with which annotated transcriptomic datasets from DNA microarray and RNA-seq experiments can be analyzed for gene expression or patient survival. Bc-GenExMiner uses Welch’s and Dunnett–Tukey–Kramer’s tests to calculate the statistical significance of gene expression differences among different subtypes [12] (https://bcgenex.ico.unicancer.fr/). The univariate Cox proportional hazards model was used to estimate the significance of prognostic values of *CLPB* expression [13]. We selected “optimal” as the splitting criterion for discretisation in our prognostic analyses.

To determine the subtype of a breast cancer patient, an approach called SSP (single sample predictor) is applied. This approach is based on comparing the average gene expression profile of the five molecular subtypes with that of the patient [18]. There are three different SSPs to classify breast cancers into intrinsic subtypes: PAM50 [19], Sorlie’s [20], and Hu’s [21]. Bc-GenExMiner can subtype breast cancer patients based on those three SSPs. *CLPB* somatic mutations in breast cancer patients were documented from Sanger COSMIC database (v101) [22] (https://cancer.sanger.ac.uk/cosmic).

### Combined bisulfite restriction analysis (COBRA)

Genomic DNAs from cell lines were isolated by DNA isolation kit (Cat. No: 69504; Qiagen, Germany) and treated with bisulfite (EpiJET Bisulfite Conversion Kit, Cat. No: K146; Thermo Sci., MA, USA). A genomic region in the CpG island, ∼150 bp downstream of the core promoter, was amplified by nested PCR with a Taq DNA polymerase (Cat. No: LSG-EP0402; Thermo Sci., MA, USA). The information about primers was given in **Supplementary Information Table 1**. The core promoter region of *CLPB* is shown by “The Eukaryotic Promoter Database” [23] (https://epd.expasy.org/epd). The amplicons (204 bp) were digested with TaqI (T↓CGA) restriction enzyme (Cat. No: ER0671; Thermo Sci., MA, USA) [24]. The expected sizes of the digestion products are 130, 48, and 26 bp. Genomic DNA of the MCF 10A cell line was in-vitro methylated by M.SssI (CpG methyltransferase) enzyme (Cat. No: EM0821) (Thermo Scientific, MA, USA) and used as a positive control for COBRA [25].

## Results

### *CLPB* expression is significantly higher in tumor samples compared to normals

*CLPB* transcript levels were analyzed with UALCAN [10] and bc-GenExMiner (v5.2) [12] in-silico tools. *CLPB* expression is significantly higher in tumors compared to normals based on TCGA datasets **(Fig. 1a, b)**. *CLPB* expression is significantly higher in all breast cancer subtypes, classified with PAM50 subtyping. But the only exception was that no significant difference was found between luminal A samples and normals based on DNA microarray data **(Fig. 1c, d)**. According to Sorlie’s and Hu’s subtyping, *CLPB* expression is significantly higher in all subtypes than in normals, based on both DNA microarray and RNA-seq datasets **(Supplementary Information Fig. 2, 3)**. The information about the datasets used to analyze the *CLPB* expression was listed in the **Supplementary Information file**.

**Fig. 1.**
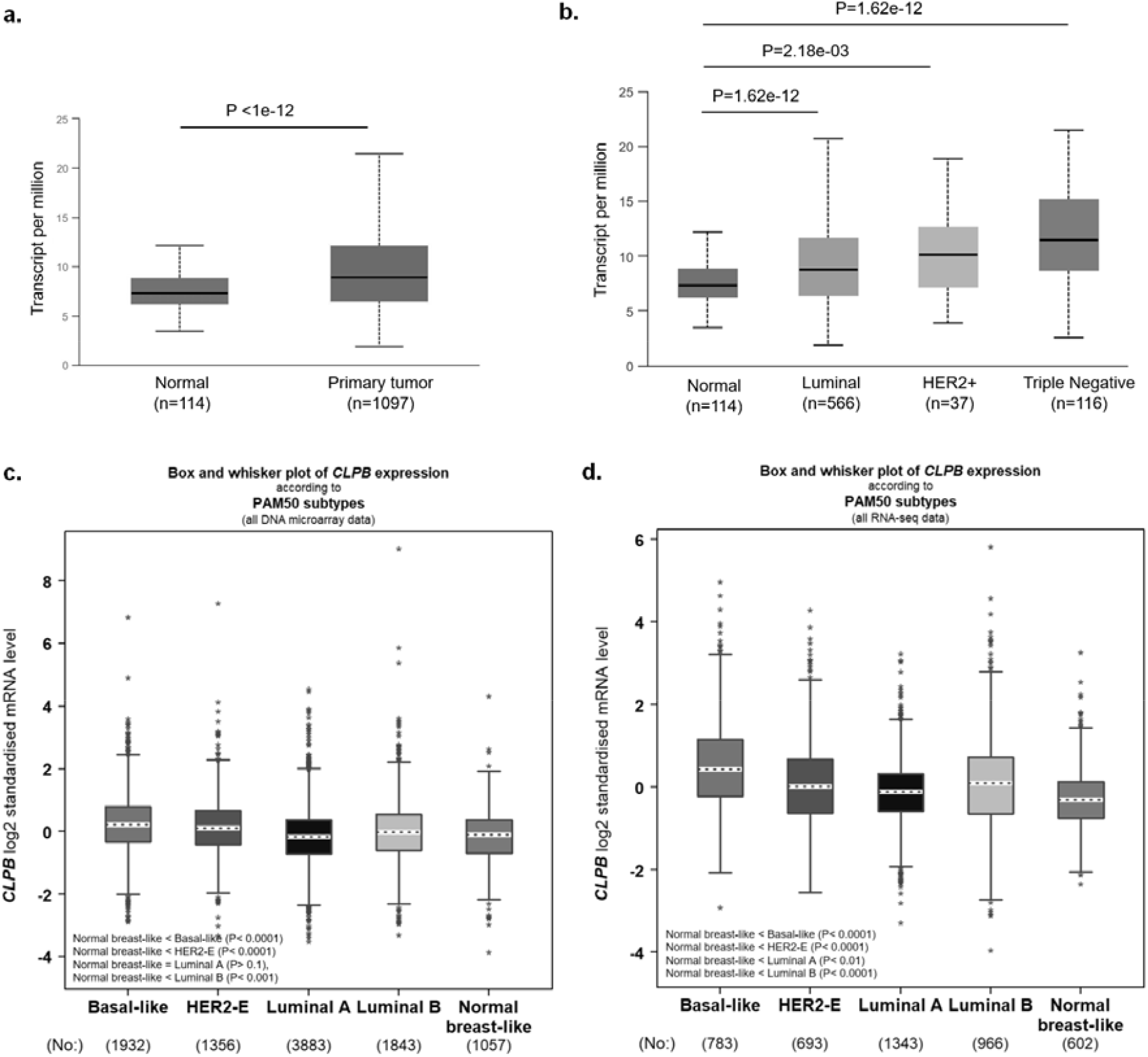
*CLPB* mRNA expression levels in clinical breast samples. UALCAN **(a, b)** and bc-GenExMiner **(c, d)** in-silico tools were used to analyze DNA microarray and RNA-seq datasets.

### High *CLPB* expression is associated with worse survival values in breast cancer

Overall survival (OS), disease-free survival (DFS), and distant metastasis-free survival (DMFS) values of breast cancer patients were evaluated with bc-GenExMiner (v5.2) [13] by employing DNA microarray and RNA-seq datasets **(Fig. 2a, b) (Supplementary Information file)**. High *CLPB* expression was found to be significantly associated with lower survival probabilities in OS, DFS, and DMFS, except for DMFS based on RNA-seq (TCGA) data.

**Fig. 2.**
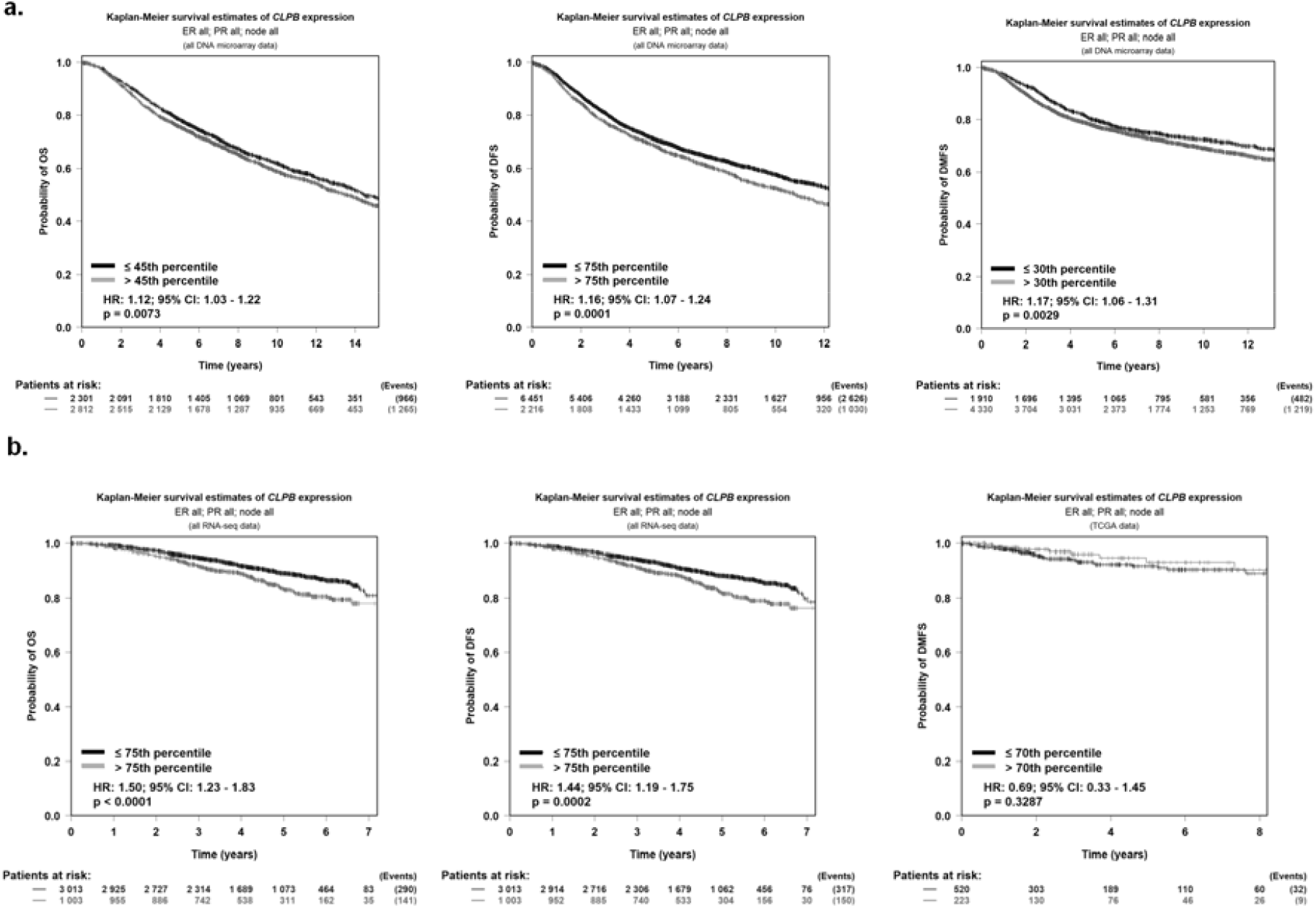
Survival statistics of breast cancer patients based on *CLPB* expression. DNA microarray **(a)** and RNA-seq **(b)** datasets were used to evaluate the survival probabilities of breast cancer patients based on their *CLPB* expression levels. Kaplan-Meier plots were generated by bc-GenExMiner (v5.2). HR: Hazard ratio; OS: Overall survival; DFS: disease-free survival; DMFS: distant metastasis-free survival.

Survival probabilities of breast cancer patients with various subtypes were also examined comprehensively **(Supplementary Information Table 2, 3, 4, 5**). According to the analysis of DNA microarray datasets in all SSPs, high *CLPB* expression in the basal-like (TNBC) subtype is unfavorable for OS and DFS but favorable for DMFS. On the contrary, *CLPB* expression negatively affects DMFS values in breast cancer samples identified as “triple negative” by immunohistochemical analysis **(Supplementary Information Fig. 4, 5)**. High *CLPB* expression is also associated with lower survival probabilities in luminal A subtype for OS (in Hu’s), DFS (in all SSPs), and DMFS (in Sorlie’s), according to the analysis of both DNA microarray and RNA-seq datasets. HER2-positive patients, subtyped based on Sorlie’s, have lower survival probabilities for OS, DFS, and DMFS.

### *CLPB* is differentially expressed in breast cancer cell lines

*CLPB* expression was analyzed in breast cancer cell lines by Multiplex Semi-quantitative RT-PCR and Q-RT-PCR. All cell lines express *CLPB* gene at moderate to high levels **(Fig. 3a, b)**.

**Fig. 3.**
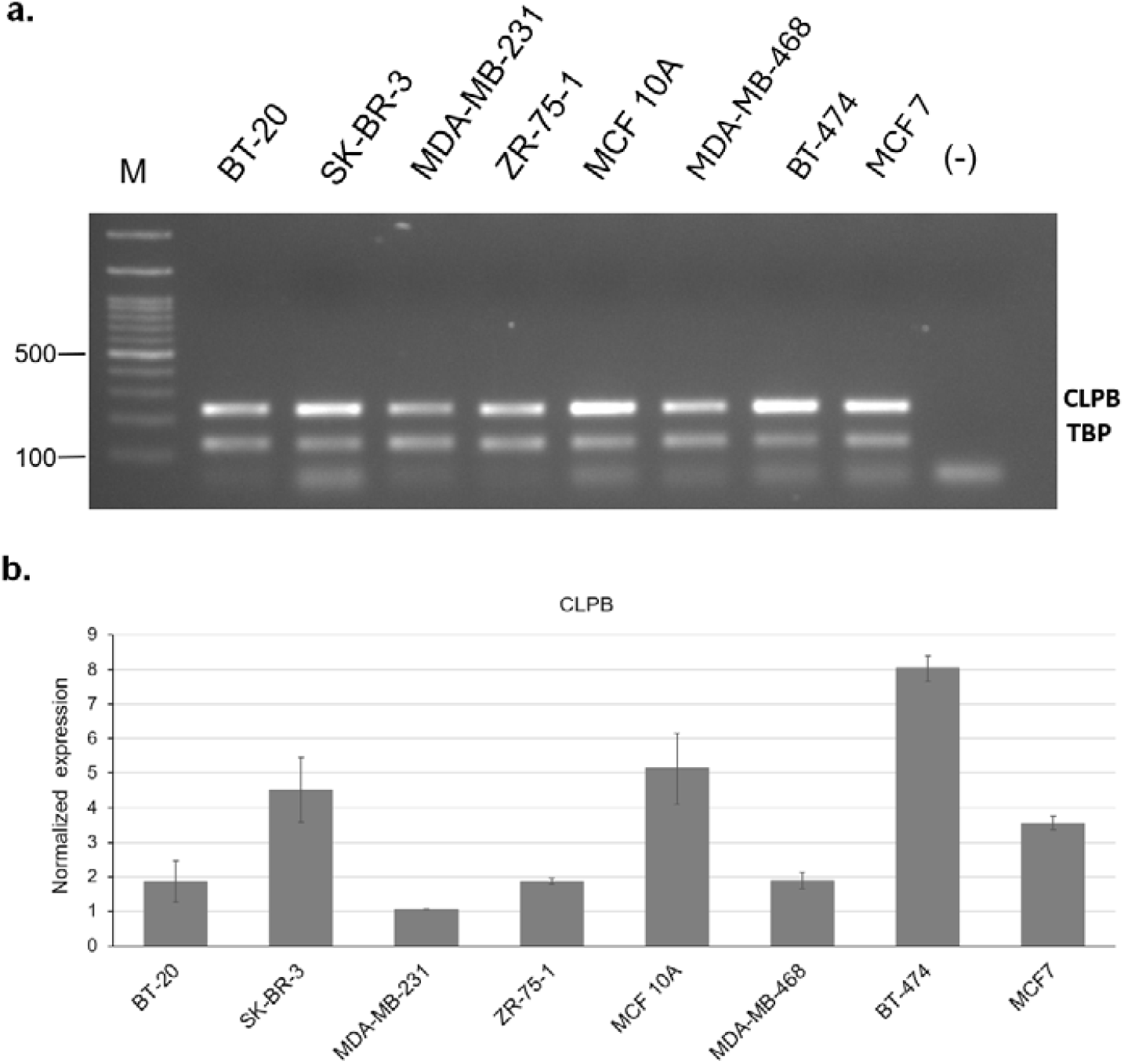
*CLPB* mRNA expression in breast cell lines. Multiplex Semi-quantitative PCR **(a)** and Q-RT-PCR **(b)** results show that *CLPB* is differentially expressed in breast cell lines (M): DNA Ladder (bp), (-): non-template PCR control. *TBP* was used for normalization. SEM (standard error of the mean) values from the three technical replicates are shown.

### *CLPB* promoter region is hypomethylated in breast cancer samples and cell lines

According to the UCSC genome browser, the core promoter region and the first exon of *CLPB* gene overlap a CpG island **(Fig. 4a)** [26] (http://genome.ucsc.edu). Based on the analysis of TCGA Illumina bead chip data by UALCAN [11], *CLPB* promoter region is hypomethylated in normal samples and all breast cancer subtypes (Beta-value = 0.3-0.25) **(Fig. 4a, b)**. But it should be noted that tumors **(Fig. 4b)** and samples of all subtypes **(Fig. 4c)**, have significantly low methylation levels compared to normals. According to our COBRA result, *CLPB* promoter region is also hypomethylated in breast cancer cell lines **(Fig. 4d)**.

**Fig. 4.**
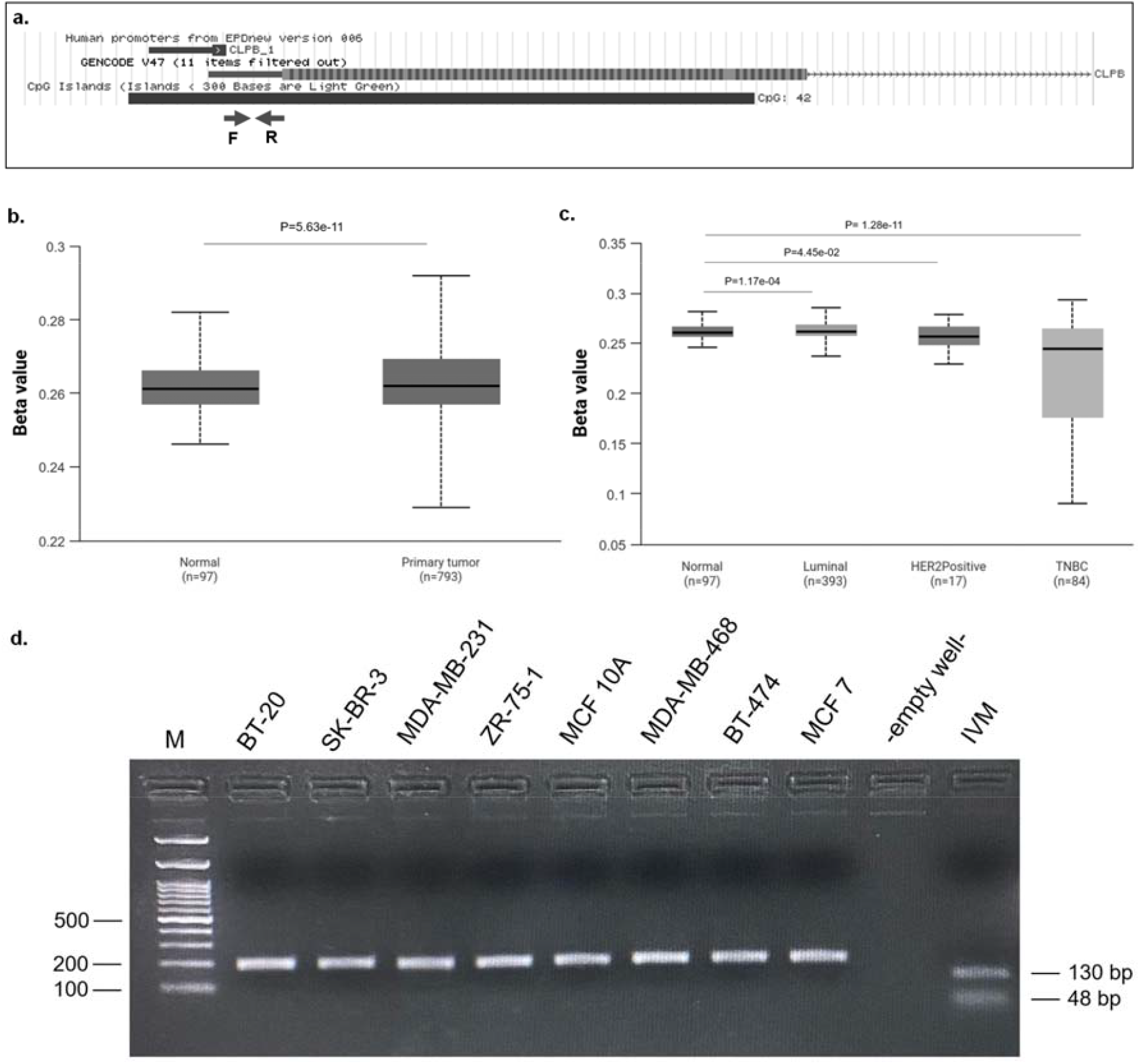
*CLPB* promoter region is hypomethylated in breast cancer. **(a)** The image taken from the UCSC genome browser (genome.ucsc.edu) and shows the core promoter region, first exon of *CLPB* gene, and CpG island from top to bottom. The approximate locations of the primers (F and R) used for COBRA are indicated by arrows. **(b, c)** *CLPB* promoter region is hypomethylated in tumor samples and samples of breast cancer subtypes **(d)** COBRA result shows that *CLPB* promoter region is hypomethylated in breast cancer cell lines. IVM (in-vitro methylated DNA) sample was used as a positive control and shows the expected digestion products; 130 bp and 48 bp. M; DNA ladder (bp).

### Genetic alterations of *CLPB* are rare in breast cancer patients

Based on the information on Sanger COSMIC database (v101) [22], the ratio of the *CLPB* somatic mutations in breast cancer patients is low (3.13%). Copy Number Variations (CNVs), either gain (2.08%) or loss (0.2%), are also rare (**Table 1**).

**Table 1.**
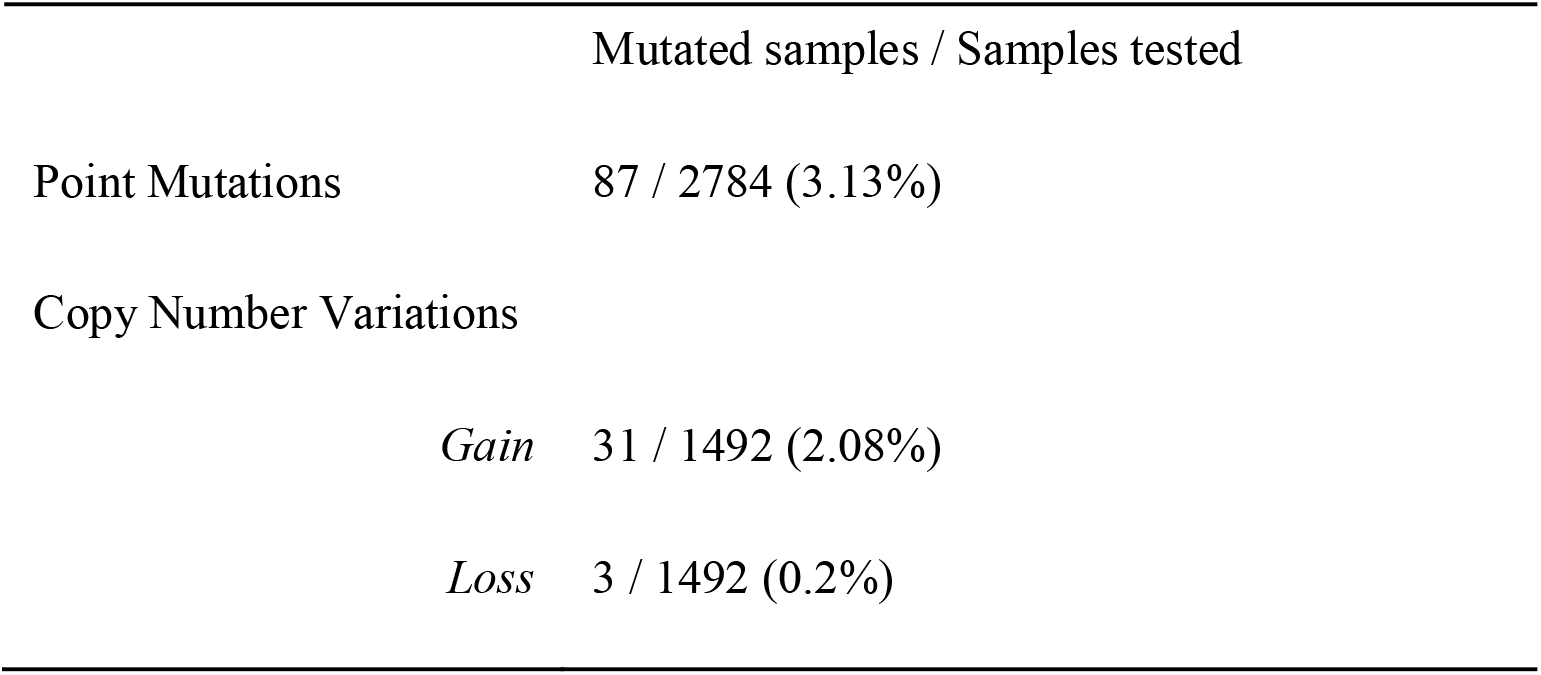
*CLPB* mutations and copy number variations in breast cancer based on the information on Sanger COSMIC database (v101)

## Discussion

CLPB mitochondrial chaperone protein has carcinogenic and anti-apoptotic functions, which have been shown in leukemia and melanoma cells [7, 8]. *CLPB* expression has also been associated with worse survival values in prostate cancer [9]. Our in-silico analysis revealed that *CLPB* expression is significantly high in stomach, colon, head and neck, esophageal, liver, and lung cancers compared to normal samples **(Supplementary Information Fig. 1)**. This background information led us to think that CLPB protein may also have a role in breast carcinogenesis and/or prognosis.

Although we have not come across a report that has studied the relationship between *CLPB* and breast cancer, a mutation in the *CLPB* gene (rs150343959, p.Arg628Cys) has been associated with early menopause in women, and CLPB protein-truncating variants have been found more frequently in women with primary ovarian insufficiency (POI) [27]. As it is known, early menopause and/or POI are factors that increase the risk of breast cancer in women [28].

In this study, expression, and survival statistics of *CLPB* gene were analyzed by employing UALCAN [10, 11] and bc-GenExMiner [12, 13] in-silico tools. Genetic alterations and promoter methylation profile that may affect *CLPB* gene expression and/or function have also been documented in breast cancer samples and cell lines. Our findings are consistent with the oncogenic character of the *CLPB* gene previously demonstrated in melanoma, leukemia, and prostate cancers. *CLPB* expression is significantly higher in breast cancer samples compared to normals, and this is true for all breast cancer subtypes. Besides, high *CLPB* expression is associated with worse overall survival (OS), disease-free survival (DFS), and distant metastasis-free survival (DMFS) values in breast cancer patients. To achieve the most consistent results, we subtyped breast cancer patients with all three SSPs by employing bc-GenExMiner (v5.2) [13, 18]. Since PAM50 has higher clinical use and validity [29, 30], we present Sorlie’s and Hu’s SSP results as additional data. **(Supplementary Information File)**.

In samples identified as basal subtype based on transcriptome profiling, *CLPB* expression positively affects DMFS. However, in samples identified as “triple negative” by immunohistochemical analysis, *CLPB* expression negatively affects DMFS **(Supplementary Information Fig. 4, 5)**. To understand this discrepancy, the following points should be considered: Tumor purity is an important factor in gene expression analyses. Stromal cells in the tumor microenvironment can influence the gene expression profile [31, 32]. Also, samples identified as triple-negative by immunohistochemical analysis may not exhibit the transcriptome profile of the basal type. One study determined that only 57% of triple-negative samples were basal-like [33]. Therefore, treatment strategies of those patients may influence DMFS data [34]. Another important issue is that TNBC is heterogeneous and has its own subtypes; LAR (luminal androgen receptor), MLIA (mesenchymal-like immune-altered), BLIA (basal-like immune-activated), and BLIS (basal-like immune-suppressed) [32, 35]. Within these subgroups, there may be differences in *CLPB* expression or differences in the proportion of tumor-infiltrating lymphocytes (TILs). High TIL levels are significantly associated with reduced distant recurrence rates in TNBC [36] and may therefore influence DMFS rates.

We used eight breast cell lines representing triple negative (basal-like) (BT-20, MDA-MB-231, MDA-MB-468), luminal A (ZR-75-1, MCF7), luminal B (BT-474), HER2-positive (SK-BR-3) breast cancer subtypes and a non-carcinogenic cell line (MCF 10A) to study the relationship between promoter methylation and *CLPB* expression. We observed moderate to high *CLPB* expression in all cell lines. However, differences in gene expression levels among cell lines representing different subtypes did not appear to be similar to those among subtypes of tumor samples. It has been discussed in several studies that the gene expression profiles of breast cell lines may not be fully representative of the expression profiles of the breast cancer subtypes they represent. Various reasons have been suggested for this situation. First of all, new mutations and/or CNVs may arise during years of cultivation that may affect gene expression profiles [37]. Most cell lines are derived from metastatic tumors and pleural effusions, and they are the most aggressive variants that could be adapted to the cell culture. Additionally, cells may lose their epithelial characteristics and acquire more mesenchymal characteristics, which leads to transcriptional distancing, particularly between ER+/luminal cell lines and the tumors they represent. Finally, the cell culture environment is significantly different from the breast cancer microenvironment. The absence of stromal and immune cells in the culture environment has been suggested as one of the major reasons for the differences in expression between cell lines and the tumor subtypes they represent. It has been stated that basal cell lines have a higher representative ability than ER+/luminal cell lines in terms of gene expression profiles [38].

It is known that hypomethylated promoter region of oncogenes could result in upregulation [39, 40]. Based on our analysis, the promoter region of *CLPB* gene is hypomethylated in all breast cancer subtypes. We validated this result in breast cancer cell lines.

In conclusion, high *CLPB* expression and its unfavorable survival outcomes in breast cancer are consistent with the oncogenic role of *CLPB* previously shown in melanoma, leukemia, and prostate cancers. The rarity of somatic mutations and CNVs in breast cancer suggests that genetic alterations do not affect gene expression or function. However, *CLPB* promoter hypomethylation would be one of the reasons for its high expression in breast cancer.

*CLPB* gene is a potential therapeutic target and biomarker candidate in breast cancer. The anti-apoptotic, cell growth/migration promoting properties of CLPB may cause resistance to chemotherapy and/or progression of breast cancer, thereby negatively affecting patient survival rates. Therefore, we are also investigating the functional effect of CLPB on breast carcinogenesis and/or chemotherapy in vitro.

## Acknowledgements

We thank Dr. I. Yulug (Bilkent University, Ankara, Turkiye) and Dr. B. Gur Dedeoglu (Ankara University, Ankara, Turkiye) for providing the cell lines.

## Author contributions

Conceptualization: Tolga Acun; Methodology: Tolga Acun; Formal analysis and investigation: Tolga Acun, Sena Sonmez, Senem Noyan; Writing - original draft preparation: Tolga Acun; Supervision: Tolga Acun.

## Data availability

The primer sequences and the information of datasets used in our study are provided in the supplementary information file.

## Ethics, Consent to Participate, and Consent to Publish declarations

Not applicable.

