## Supplementary Information for "Hypomethylation may drive *CLPB* overexpression which predicts poor survival outcomes in breast cancer"

**Journal name:** BioRxiv

**Author name:** Sena Sonmez[a], Senem Noyan[b], Tolga Acun[a*]

[a] Department of Molecular Biology and Genetics, Faculty of Science, Zonguldak Bulent Ecevit University, Zonguldak, 67100, Turkiye.

[b] Biotechnology Institute, Ankara University, Ankara, Turkiye.

* Corresponding author

Tolga Acun, Ph.D.

Department of Molecular Biology and Genetics, Faculty of Science, Zonguldak Bulent Ecevit University

Zonguldak, 67100, Turkiye

**Supp. Info. Table 1.** Primers

| **Semi-Q-RT-PCR and Q-RT-PCR** | | |
|---|---|---|
| **Primer Name** | **Sequence (5'-3')** | **Product Size (bp)** |
| CLPB-QPCR-F | GGAGATGAGCCGTAACCGTA | 241 |
| CLPB-QPCR-R | CCTTTGCTTGGCTCTCTTGG | |
| TBP-F* | TGCACAGGAGCCAAGAGTGAA | 132 |
| TBP-R* | CACATCACAGCTCCCCACCA | |
| **COBRA** | | |
| **Primer Name** | **Sequence (5'-3')** | **Product Size (bp)** |
| CLPB-COBRA-F | TTTTTGGTGTTGAGGAGAAAAGTAT | 379 |
| CLPB-COBRA-R | TCTTACTATAACAATAAACCACCAAC | |
| CLPB-COBRA-NF | GGAATGTGATTATTGGGAGTTT | 204 |
| CLPB-COBRA-NR | TATCCTATCCTAAAAATATTTCTTC | |

* Gur-Dedeoglu B. *et al.* (2009)

***CLPB* Ensembl Gene ID:** ENSG00000162129

**Supp. Info. Table 2** Survival values of *CLPB* expression in breast cancer subtypes (PAM50) examined by bc-GenExMiner (v5.2) in-silico tool.

| | Overall Survival | | Disease-Free Survival | | Distant Metastasis-Free Survival | |
|---|---|---|---|---|---|---|
| | DNA Microarray | RNA-seq | DNA Microarray | RNA-seq | DNA Microarray | RNA-seq (TCGA) |
| Basal-like | **Unfavorable**<br>P= 0.04<br>HR = 1.29<br>n = 853 | N.S.<br>P = 0.15<br>HR = 1.35<br>n = 712 | **Unfavorable**<br>P= 0.004<br>HR = 1.35<br>n = 1599 | N.S.<br>P = 0.22<br>HR = 1.28<br>n = 712 | **Favorable**<br>P= 0.04<br>HR = 0.78<br>n = 1073 | N.S.<br>P = 0.13<br>HR = 0.43<br>n = 167 |
| Luminal A | N.S.<br>P = 0.16<br>HR = 1.10<br>n = 2309 | **Unfavorable**<br>P = 0.01<br>HR = 2.32<br>n = 1205 | **Unfavorable**<br>P = 0.001<br>HR = 1.22<br>n = 3355 | **Unfavorable**<br>P = 0.007<br>HR = 1.93<br>n = 1205 | N.S.<br>P= 0.09<br>HR = 1.21<br>n = 2654 | N.S.<br>P = 0.13<br>HR = 3.35<br>n = 255 |
| Luminal B | **Favorable**<br>P = 0.03<br>HR = 0.78<br>n = 786 | N.S.<br>P = 0.23<br>HR = 1.28<br>n = 891 | N.S.<br>P = 0.13<br>HR = 0.87<br>n = 1531 | N.S.<br>P = 0.28<br>HR = 1.24<br>n = 891 | N.S.<br>P = 0.19<br>HR = 0.86<br>n = 1068 | N.S.<br>P = 0.19<br>HR = 2.45<br>n = 141 |
| HER2 Positive | N.S.<br>P = 0.25<br>HR = 1.18<br>n = 608 | N.S.<br>P = 0.35<br>HR = 1.25<br>n = 619 | N.S.<br>P = 0.13<br>HR = 1.16<br>n = 1128 | N.S.<br>P = 0.44<br>HR = 1.24<br>n = 619 | N.S.<br>P = 0.07<br>HR = 1.25<br>n = 743 | N.S.<br>P = 0.23<br>HR = 0.30<br>n = 112 |

N.S.: Not Significant
HR: Hazard ratio
Favorable: High expression is associated with high survival probability
Unfavorable: High expression is associated with low survival probability

**Supp. Info. Table 3** Survival values of *CLPB* expression in breast cancer subtypes (Sorlie's) examined by bc-GenExMiner (v5.2) in-silico tool.

| | Overall Survival | | Disease-Free Survival | | Distant Metastasis-Free Survival | |
|---|---|---|---|---|---|---|
| | DNA Microarray | RNA-seq | DNA Microarray | RNA-seq | DNA Microarray | RNA-seq (TCGA) |
| Basal-like | **Unfavorable**<br>P= 0.04<br>HR = 1.35<br>n = 585 | N.S.<br>P = 0.30<br>HR = 1.29<br>n = 521 | **Unfavorable**<br>P= 0.01<br>HR = 1.34<br>n = 1182 | N.S.<br>P = 0.33<br>HR = 1.26<br>n = 521 | **Favorable**<br>P= 0.009<br>HR = 0.69<br>n = 852 | N.S.<br>P = 0.17<br>HR = 0.46<br>n = 151 |
| Luminal A | N.S.<br>P = 0.13<br>HR = 1.15<br>n = 1117 | N.S.<br>P = 0.057<br>HR = 1.67<br>n = 1359 | **Unfavorable**<br>P = 0.004<br>HR = 1.27<br>n = 2274 | **Unfavorable**<br>P = 0.01<br>HR = 1.54<br>n = 1359 | **Unfavorable**<br>P= 0.03<br>HR = 1.31<br>n = 1643 | **Unfavorable**<br>P = 0.009<br>HR = 4.99<br>n = 270 |
| Luminal B | N.S.<br>P = 0.5<br>HR = 0.95<br>n = 1394 | N.S.<br>P = 0.25<br>HR = 0.74<br>n = 571 | N.S.<br>P = 0.37<br>HR = 1.07<br>n = 1904 | N.S.<br>P = 0.28<br>HR = 0.76<br>n = 571 | N.S.<br>P = 0.45<br>HR = 1.08<br>n = 1660 | N.S.<br>P = 0.052<br>HR = 0.11<br>n = 93 |
| HER2 Positive | N.S.<br>P = 0.27<br>HR = 1.30<br>n = 337 | **Unfavorable**<br>P = 0.008<br>HR = 1.79<br>n = 548 | **Unfavorable**<br>P = 0.03<br>HR = 1.30<br>n = 781 | **Unfavorable**<br>P = 0.006<br>HR = 1.80<br>n = 548 | **Unfavorable**<br>P = 0.03<br>HR = 1.39<br>n = 507 | N.S.<br>P = 0.07<br>HR = 0.14<br>n = 87 |

N.S.: Not Significant
HR: Hazard ratio
Favorable: High expression is associated with high survival probability
Unfavorable: High expression is associated with low survival probability

**Supp. Info. Table 4** Survival values of *CLPB* expression in breast cancer subtypes (Hu's) examined by bc-GenExMiner (v5.2) in-silico tool.

| | Overall Survival | | Disease-Free Survival | | Distant Metastasis-Free Survival | |
|---|---|---|---|---|---|---|
| | DNA Microarray | RNA-seq | DNA Microarray | RNA-seq | DNA Microarray | RNA-seq (TCGA) |
| Basal-like | **Unfavorable**<br>P= 0.02<br>HR = 1.33<br>n = 836 | N.S.<br>P = 0.20<br>HR = 1.25<br>n = 864 | **Unfavorable**<br>P= 0.01<br>HR = 1.28<br>n = 1713 | N.S.<br>P = 0.32<br>HR = 1.19<br>n = 864 | **Favorable**<br>P= 0.006<br>HR = 0.72<br>n = 1161 | N.S.<br>P = 0.12<br>HR = 0.41<br>n = 187 |
| Luminal A | **Unfavorable**<br>P = 0.01<br>HR = 1.19<br>n = 1833 | **Unfavorable**<br>P = 0.01<br>HR = 1.75<br>n = 1022 | **Unfavorable**<br>P = 0.001<br>HR = 1.26<br>n = 2740 | **Unfavorable**<br>P = 0.007<br>HR = 1.95<br>n = 1022 | N.S.<br>P= 0.12<br>HR = 1.19<br>n = 2167 | N.S.<br>P = 0.08<br>HR = 3.81<br>n = 202 |
| Luminal B | N.S.<br>P = 0.18<br>HR = 0.85<br>n = 634 | N.S.<br>P = 0.32<br>HR = 1.23<br>n = 858 | N.S.<br>P = 0.27<br>HR = 0.91<br>n = 1362 | N.S.<br>P = 0.19<br>HR = 1.30<br>n = 858 | N.S.<br>P = 0.18<br>HR = 1.20<br>n = 919 | N.S.<br>P = 0.051<br>HR = 4.91<br>n = 152 |
| HER2 Positive | N.S.<br>P = 0.10<br>HR = 0.78<br>n = 390 | N.S.<br>P = 0.08<br>HR = 1.91<br>n = 347 | N.S.<br>P = 0.24<br>HR = 1.18<br>n = 704 | N.S.<br>P = 0.15<br>HR = 1.62<br>n = 347 | N.S.<br>P = 0.29<br>HR = 1.24<br>n = 455 | N.S.<br>P = 0.08<br>HR = 0.21<br>n = 64 |

N.S.: Not Significant
HR: Hazard ratio
Favorable: High expression is associated with high survival probability
Unfavorable: High expression is associated with low survival probability

**Supp. Info. Table 5** Survival values of *CLPB* expression in breast cancer subtypes (RSSPC) examined by bc-GenExMiner (v5.2) in-silico tool.

| | Overall Survival | | Disease-Free Survival | | Distant Metastasis-Free Survival | |
|---|---|---|---|---|---|---|
| | DNA Microarray | RNA-seq | DNA Microarray | RNA-seq | DNA Microarray | RNA-seq (TCGA) |
| Basal-like | **Unfavorable**<br>P= 0.04<br>HR = 1.36<br>n = 522 | N.S.<br>P = 0.29<br>HR = 1.30<br>n = 488 | **Unfavorable**<br>P= 0.01<br>HR = 1.40<br>n = 1064 | N.S.<br>P = 0.29<br>HR = 1.30<br>n = 488 | **Favorable**<br>P= 0.008<br>HR = 0.67<br>n = 756 | N.S.<br>P = 0.23<br>HR = 0.49<br>n = 145 |
| Luminal A | N.S.<br>P = 0.26<br>HR = 0.86<br>n = 736 | **Unfavorable**<br>P = 0.03<br>HR = 2.16<br>n = 633 | **Unfavorable**<br>P = 0.03<br>HR = 1.24<br>n = 1302 | **Unfavorable**<br>P = 0.01<br>HR = 2.17<br>n = 633 | N.S.<br>P= 0.23<br>HR = 1.21<br>n = 973 | N.S.<br>P = 0.20<br>HR = 3.55<br>n = 144 |
| Luminal B | N.S.<br>P = 0.39<br>HR = 0.78<br>n = 155 | N.S.<br>P = 0.40<br>HR = 1.44<br>n = 194 | N.S.<br>P = 0.08<br>HR = 0.75<br>n = 332 | N.S.<br>P = 0.43<br>HR = 1.40<br>n = 194 | N.S.<br>P = 0.13<br>HR = 0.71<br>n = 246 | N.S.<br>P = 0.63<br>HR = 0.51<br>n = 31 |
| HER2 Positive | N.S.<br>P = 0.15<br>HR = 0.63<br>n = 120 | N.S.<br>P = 0.20<br>HR = 2.16<br>n = 172 | N.S.<br>P = 0.09<br>HR = 1.42<br>n = 263 | N.S.<br>P = 0.14<br>HR = 2.42<br>n = 172 | N.S.<br>P = 0.19<br>HR = 1.44<br>n = 157 | N.S.<br>P = 0.13<br>HR = 0.18<br>n = 42 |

RSSPC: Robust SSP classification based on patients classified in the same subtype with the three SSPs (Sorlie, Hu, PAM50)
N.S.: Not Significant
HR: Hazard ratio
Favorable: High expression is associated with high survival probability
Unfavorable: High expression is associated with low survival probability

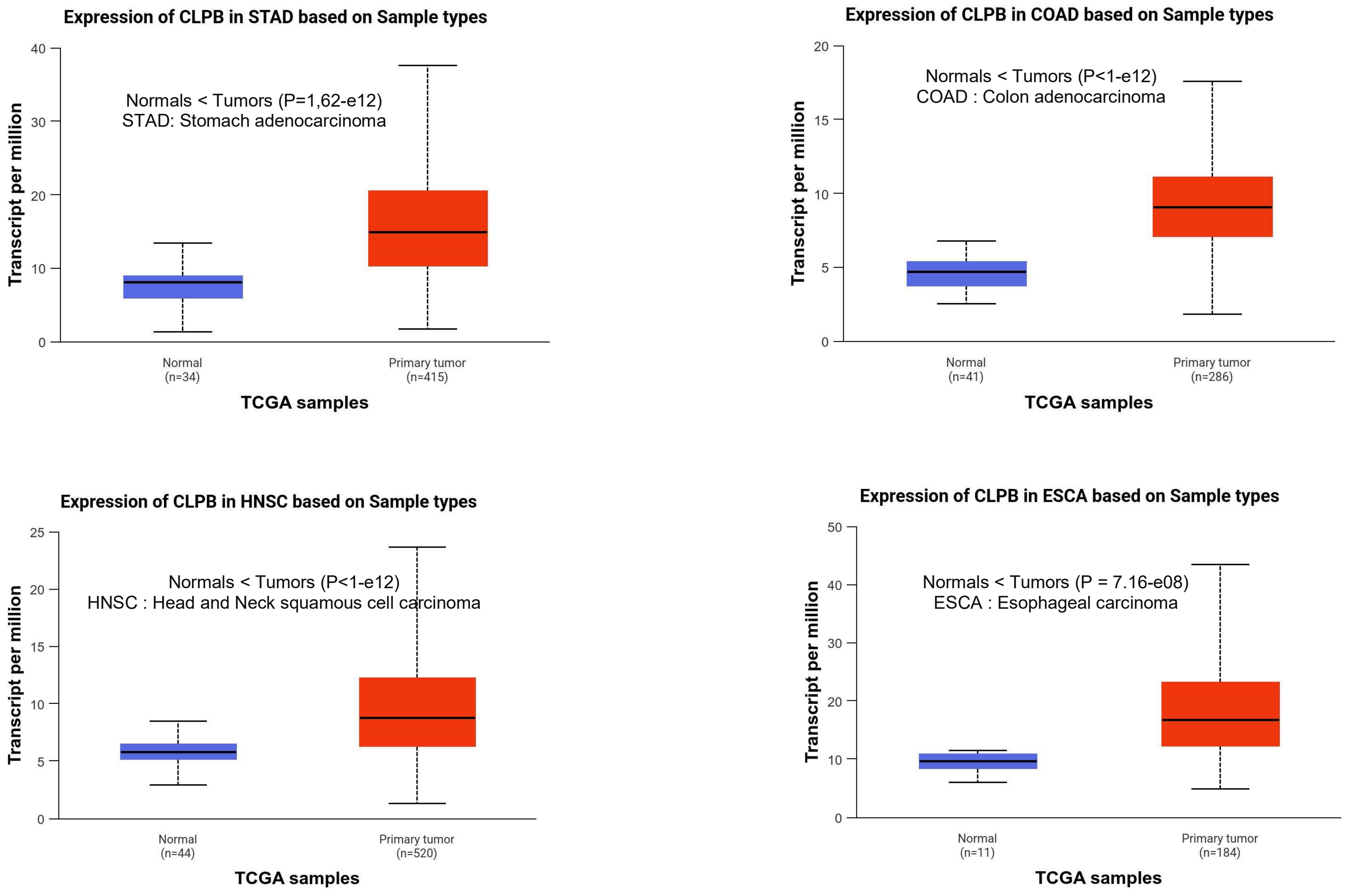


**Supp. Info. Fig. 1** *CLPB* mRNA expression values on various cancer types based on TCGA datasets analysed by UALCAN in-silico tool.

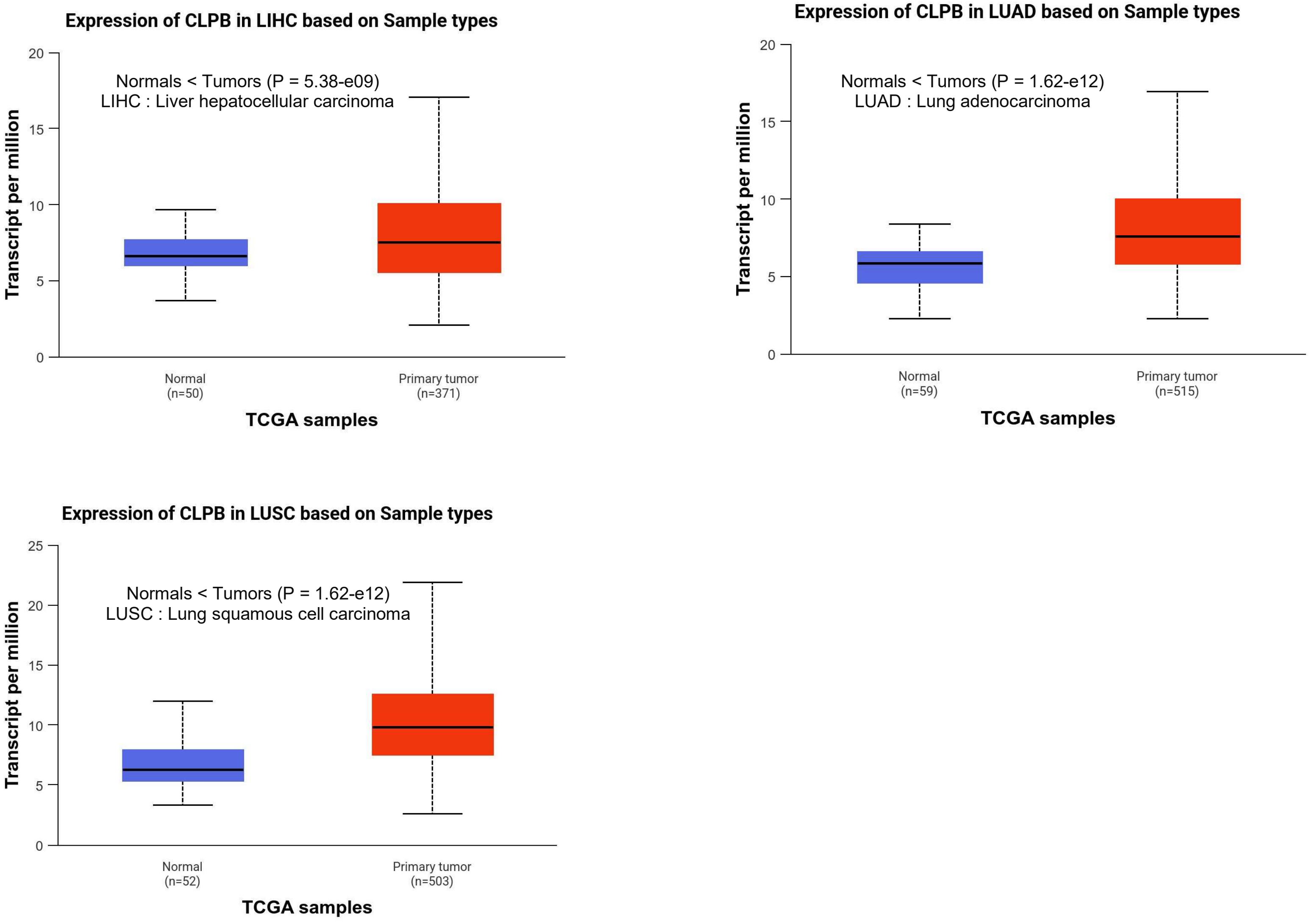


**Supp. Info. Fig. 1 (cont.)** *CLPB* mRNA expression values on various cancer types based on TCGA datasets analysed by UALCAN in-silico tool.

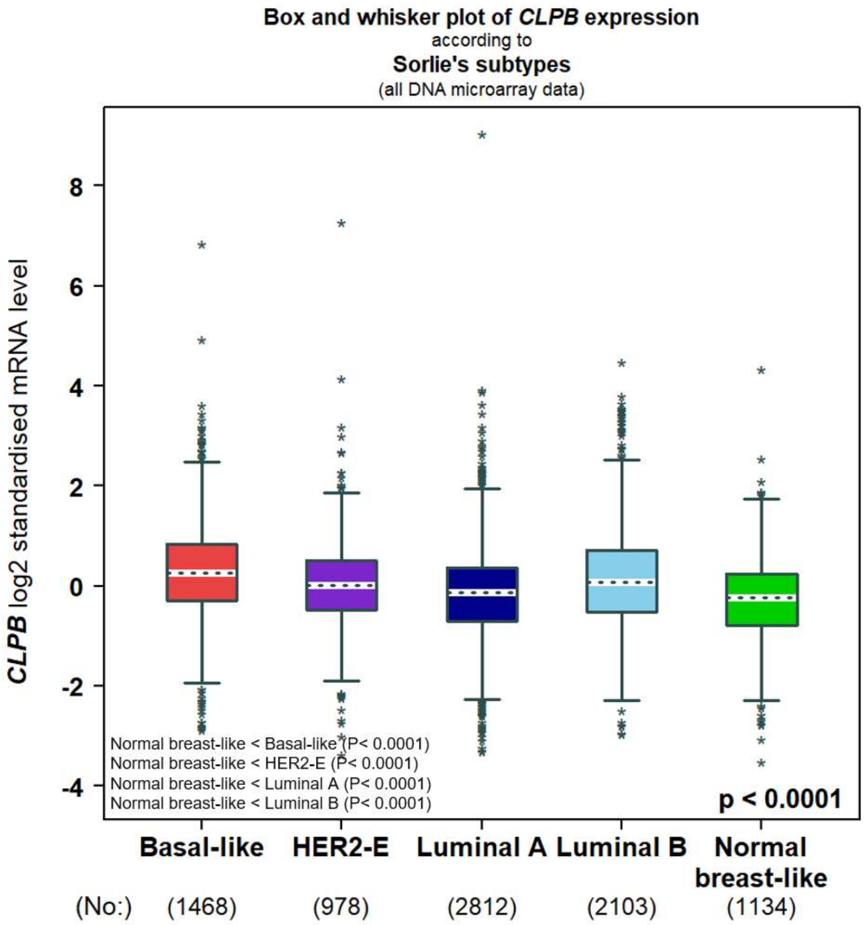


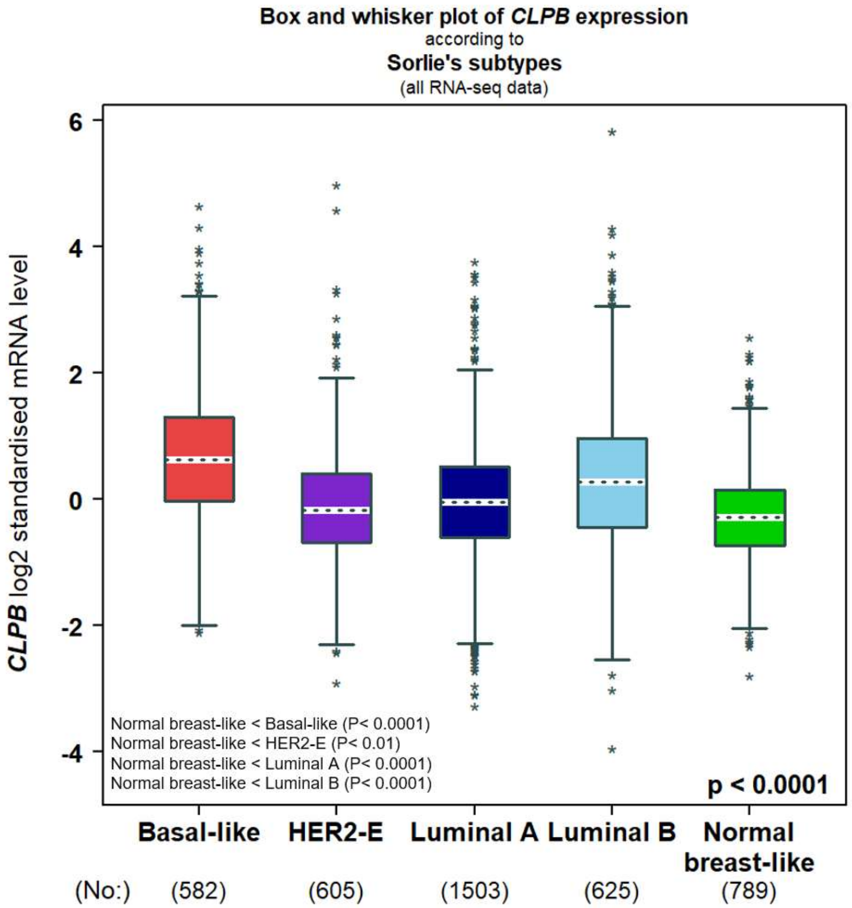


**Supp. Info. Fig. 2** *CLPB* expression is significantly higher in all subtypes compared to normals, based on analyses of DNA microarray and RNA-seq datasets. Breast cancer patients were subtyped based on Sorlie's subtyping (Sorlie et al., 2001).

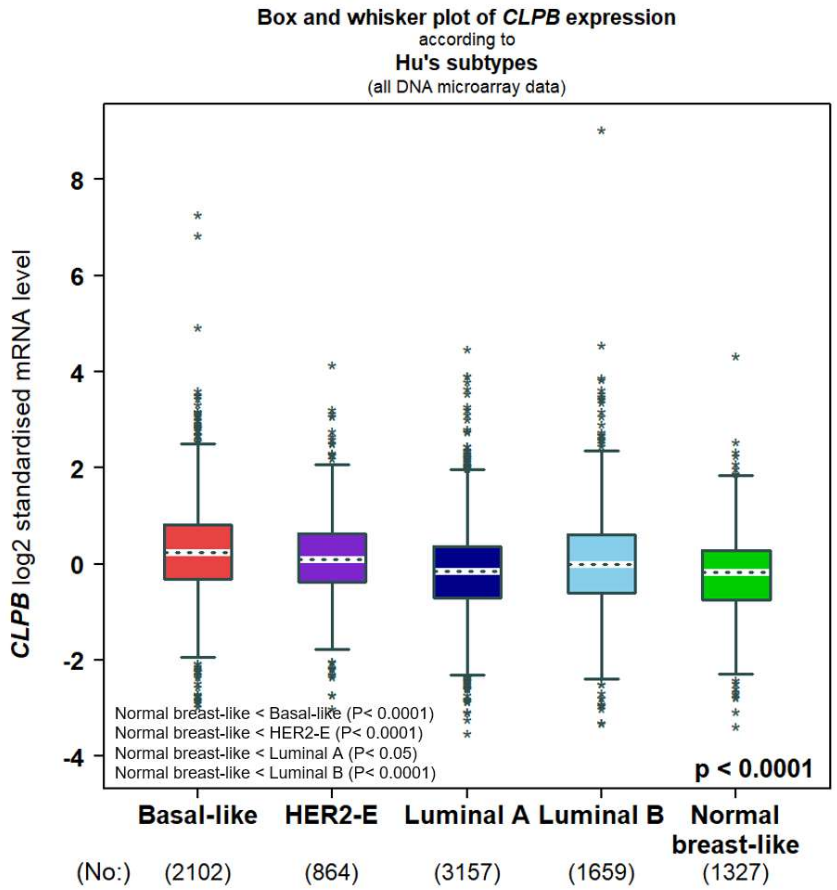


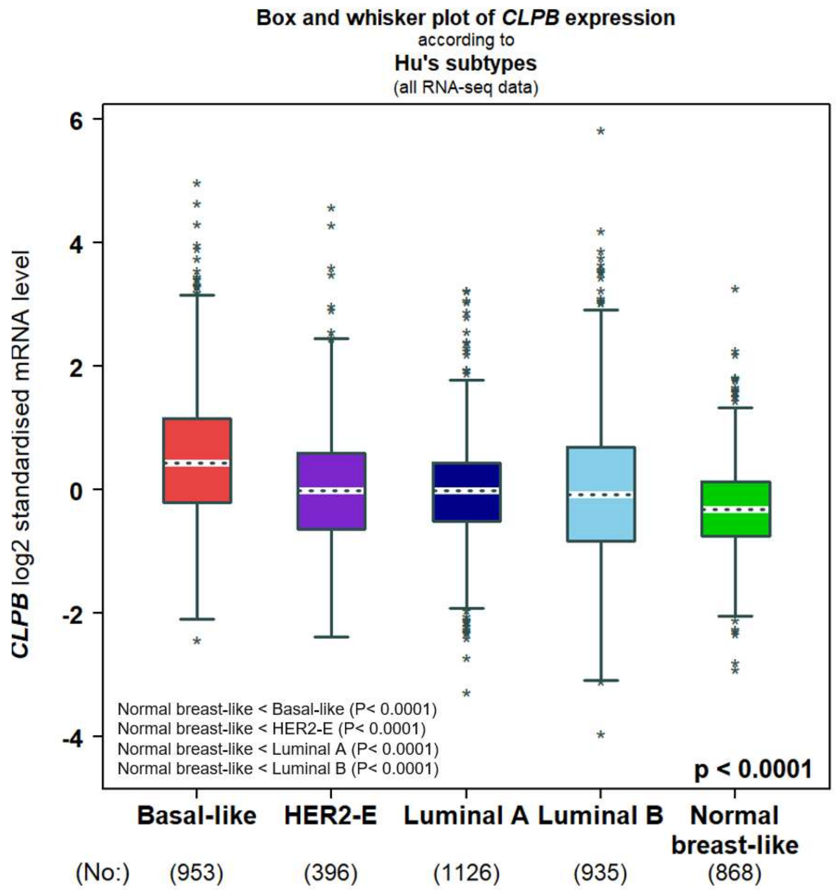


**Supp. Info. Fig. 3** *CLPB* expression is significantly higher in all subtypes compared to normals, based on analyses of DNA microarray and RNA-seq datasets. Breast cancer patients were subtyped based on Hu's subtyping (Hu et al., 2006).

| **CLPB univariate Cox analysis (TNBC (IHC) and/or basal-like (PAM50))** (all DNA microarray data - TNBC (IHC) - optimised split) | | | | | | | | | |
|---|---|---|---|---|---|---|---|---|---|
| Time-to-event endpoints | p-value | HR | 95% CI | Good prognosis' RNA level | No. patients | No. events | Kaplan-Meier curves View | Kaplan-Meier curves Save figure | |
| DFS | 0.0204 | 1.36 | 1.05 - 1.76 | ↘ | 789 | 368 | | PNG | SVG |
| DMFS | 0.0469 | 1.44 | 1.01 - 2.05 | ↘ | 546 | 181 | | PNG | SVG |
| OS | 0.0507 | 1.37 | 1.00 - 1.89 | ↘ | 463 | 249 | | PNG | SVG |

**Online Resource Fig. 4** Survival values of *CLPB* expression in TNBC samples classified by IHC examined by bc-GenExMiner (v5.2) in-silico tool.

| **CLPB univariate Cox analysis (TNBC (IHC) subtypes)** (all DNA microarray data - All TNBC subtypes - optimised split) | | | | | | | | | |
|---|---|---|---|---|---|---|---|---|---|
| Time-to-event endpoints | p-value | HR | 95% CI | Good prognosis' RNA level | No. patients | No. events | Kaplan-Meier curves View | Kaplan-Meier curves Save figure | |
| DFS | 0.0464 | 1.24 | 1.00 - 1.54 | ↘ | 866 | 350 | | PNG | SVG |
| DMFS | 0.0480 | 1.34 | 1.00 - 1.80 | ↘ | 641 | 185 | | PNG | SVG |
| OS | 0.0921 | 1.24 | 0.97 - 1.58 | ↘ | 654 | 261 | | PNG | SVG |

**Supp. Info. Fig. 5** Survival values of *CLPB* expression in TNBC subtypes examined by bc-GenExMiner (v5.2) in-silico tool.

**DNA microarray datasets which are used to analyze the *CLPB* mRNA expression by using bc-GenExMiner (v5.2)**

**Population and event data:**

| Ver | Study code | Original data | | Filtered data | | Final data |
|---|---|---|---|---|---|---|
| | | Reference | No. patients | Instrinsic molecular subtypes -selected patients | Gene found | No. patients |
| 1.0 | **Rosetta2002** | Van de Vijver *et al.*, 2002 | 295 | 295 | | 295 |
| 1.0 | **PNAS1732912100** | Sotiriou *et al.*, 2003 | 99 | 99 | | 0 |
| 1.0 | **GSE1379** | Ma *et al.*, 2004 | 59 | 59 | | 59 |
| 1.0 | **GSE2603** | Minn *et al.*, 2005 | 82 | 82 | | 82 |
| 1.0 | **GSE1456** | Pawitan *et al.*, 2005 | 159 | 159 | | 159 |
| 1.0 | **GSE2034** | Wang *et al.*, 2005 | 286 | 286 | | 286 |
| 1.0 | **GSE2741** | Weigelt *et al.*, 2005 | 50 | 50 | | 50 |
| 1.0 | **GSE3143** | Bild *et al.*, 2006 | 158 | 158 | | 0 |
| 1.0 | **E_TABM_158** | Chin *et al.*, 2006 | 112 | 112 | | 112 |
| 1.0 | **GSE4922** | Ivshina *et al.*, 2006 | 249 | 249 | | 249 |
| 1.0 | **GSE7390** | Desmedt *et al.*, 2007 | 198 | 198 | | 198 |
| 1.0 | **GSE6532** | Loi *et al.*, 2007 | 267 | 267 | | 267 |
| 1.0 | **GSE5327** | Minn *et al.*, 2007 | 58 | 58 | | 58 |
| 1.0 | **E_UCON_1** | Naderi *et al.*, 2007 | 135 | 135 | | 135 |
| 1.0 | **GSE7849** | Anders *et al.*, 2008 | 75 | 75 | | 0 |
| 1.0 | **GSE9893** | Chanrion *et al.*, 2008 | 151 | 151 | | 151 |
| 1.0 | **GSE9195** | Loi *et al.*, 2008 | 77 | 77 | | 77 |
| 1.0 | **GSE10510** | Calabrò *et al.*, 2009 | 139 | 139 | | 139 |
| 1.0 | **GSE11264** | Jézéquel *et al.*, 2009 | 252 | 0 | | 0 |
| 1.1 | **GSE11121** | Schmidt *et al.*, 2008 | 200 | 200 | | 200 |
| 1.1 | **GSE12093** | Zhang *et al.*, 2009 | 136 | 136 | | 136 |
| 3.1 | **GSE8757** | Chin *et al.*, 2007 | 171 | 171 | | 171 |
| 3.1 | **GSE7378** | Zhou *et al.*, 2007 | 54 | 54 | | 54 |
| 3.1 | **GSE16391** | Desmedt *et al.*, 2009 | 55 | 55 | | 55 |
| 3.1 | **GSE22133** | Jönsson *et al.*, 2010 | 346 | 345 | | 345 |
| 3.1 | **GSE19615** | Li *et al.*, 2010 | 115 | 115 | | 115 |
| 3.1 | **GSE17907** | Sircoulomb *et al.*, 2010 | 55 | 55 | | 55 |
| 3.1 | **GSE22219** | Buffa *et al.*, 2011 | 216 | 216 | | 216 |
| 3.1 | **GSE20711** | Dedeurwaerder *et al.*, 2011 | 85 | 85 | | 85 |
| 3.1 | **GSE26971** | Filipits *et al.*, 2011 | 277 | 277 | | 277 |
| 3.1 | **GSE25055** | Hatzis *et al.*, 2011 | 309 | 309 | | 309 |

| | | | | | | |
|---|---|---|---|---|---|---|
| 3.1 | **GSE20685** | Kao *et al.*, 2011 | 296 | 296 | | 296 |
| 3.1 | **GSE21653** | Sabatier *et al.*, 2011 | 239 | 239 | | 239 |
| 3.1 | **GSE16987** | Wang *et al.*, 2011 | 149 | 149 | | 149 |
| 3.1 | **GSE45255** | Nagalla *et al.*, 2013 | 41 | 41 | | 41 |
| 4.3 | **GSE2109** | expO *et al.*, 2005 | 298 | 298 | | 298 |
| 4.3 | **GSE8193** | Yau *et al.*, 2007 | 47 | 47 | | 47 |
| 4.3 | **GSE20462** | Parris *et al.*, 2010 | 94 | 94 | | 94 |
| 4.3 | **GSE17705** | Symmans *et al.*, 2010 | 43 | 43 | | 43 |
| 4.3 | **GSE24450** | Heikkinen *et al.*, 2011 | 174 | 174 | | 174 |
| 4.3 | **GSE31448** | Sabatier *et al.*, 2011 | 71 | 71 | | 71 |
| 4.3 | **METABRIC** | Curtis *et al.*, 2012 | 1 980 | 1 980 | | 1 980 |
| 4.3 | **E_MTAB_365** | Guedj *et al.*, 2012 | 536 | 536 | | 536 |
| 4.3 | **GSE30682** | Servant *et al.*, 2012 | 343 | 343 | | 343 |
| 4.3 | **GSE42568** | Clarke *et al.*, 2013 | 104 | 104 | | 104 |
| 4.3 | **GSE40115** | Larsen *et al.*, 2013 | 183 | 183 | | 183 |
| 4.3 | **GSE55348** | Castagnoli *et al.*, 2014 | 53 | 53 | | 53 |
| 4.3 | **GSE43358** | Fumagalli *et al.*, 2014 | 56 | 56 | | 56 |
| 4.3 | **GSE36295** | Merdad *et al.*, 2014 | 45 | 45 | | 45 |
| 4.3 | **GSE37751** | Terunuma *et al.*, 2014 | 55 | 55 | | 55 |
| 4.3 | **GSE76274** | Burstein *et al.*, 2015 | 66 | 66 | | 66 |
| 4.3 | **GSE97177** | Biermann *et al.*, 2017 | 53 | 53 | | 53 |
| 4.5 | **GSE12276** | Bos *et al.*, 2009 | 204 | 204 | | 204 |
| 4.5 | **GSE18864** | Silver *et al.*, 2010 | 75 | 75 | | 75 |
| 4.6 | **GSE80999** | Aure *et al.*, 2017 | 381 | 381 | | 381 |
| 4.6 | **GSE86166** | Prabhakaran *et al.*, 2017 | 366 | 366 | | 366 |
| | | **Total:** | **10 872** | **10 619** | **52** | **10 287** |

**Summary:**
Expression analysis for *CLPB* with all intrinsic molecular subtypes splitting criteria (n = 10 287).

**RNA-seq datasets which are used to analyze the *CLPB* mRNA expression by using bc-GenExMiner (v5.2)**

**Population and event data:**

| Ver | Study code | Original data | | Filtered data | | Final data |
|---|---|---|---|---|---|---|
| | | Reference | No. patients | Instrinsic molecular subtypes -selected patients | Gene found | No. patients |
| 4.3 | **TCGA** | TCGA *et al.*, 2012 | 743 | 743 | | 743 |
| 4.3 | **SCAN-B / GSE96058** | Saal *et al.*, 2015 | 3 273 | 3 273 | | 3 273 |
| 4.3 | **SCAN-B / GSE81538** | Brueffer *et al.*, 2018 | 405 | 405 | | 405 |
| | | **Total:** | **4 421** | **4 421** | **3** | **4 421** |

**Summary:**
Expression analysis for *CLPB* with all intrinsic molecular subtypes splitting criteria (n = 4 421).

**DNA microarray datasets which are used to analyze the overall survival rates by using bc-GenExMiner (v5.2)**

**Population and event data:**

| Ver | Study code | Original data | | Filtered data | | Final data | |
|---|---|---|---|---|---|---|---|
| | | Reference | No. patients | (ER all, PR all, N all and OS) -selected patients | Gene found | No. patients | No. OS |
| 1.0 | **Rosetta2002** | Van de Vijver *et al.*, 2002 | 295 | 295 | | 295 | 79 |
| 1.0 | **PNAS1732912100** | Sotiriou *et al.*, 2003 | 99 | 99 | | 0 | 0 |
| 1.0 | **GSE1379** | Ma *et al.*, 2004 | 59 | 0 | | 0 | 0 |
| 1.0 | **GSE2603** | Minn *et al.*, 2005 | 82 | 0 | | 0 | 0 |
| 1.0 | **GSE1456** | Pawitan *et al.*, 2005 | 159 | 159 | | 159 | 40 |
| 1.0 | **GSE2034** | Wang *et al.*, 2005 | 286 | 0 | | 0 | 0 |
| 1.0 | **GSE2741** | Weigelt *et al.*, 2005 | 50 | 50 | | 50 | 10 |
| 1.0 | **GSE3143** | Bild *et al.*, 2006 | 158 | 158 | | 0 | 0 |
| 1.0 | **E_TABM_158** | Chin *et al.*, 2006 | 112 | 112 | | 112 | 35 |
| 1.0 | **GSE4922** | Ivshina *et al.*, 2006 | 249 | 0 | | 0 | 0 |
| 1.0 | **GSE7390** | Desmedt *et al.*, 2007 | 198 | 198 | | 198 | 56 |
| 1.0 | **GSE6532** | Loi *et al.*, 2007 | 267 | 0 | | 0 | 0 |
| 1.0 | **GSE5327** | Minn *et al.*, 2007 | 58 | 0 | | 0 | 0 |
| 1.0 | **E_UCON_1** | Naderi *et al.*, 2007 | 135 | 135 | | 135 | 47 |
| 1.0 | **GSE7849** | Anders *et al.*, 2008 | 75 | 0 | | 0 | 0 |
| 1.0 | **GSE9893** | Chanrion *et al.*, 2008 | 151 | 151 | | 151 | 41 |
| 1.0 | **GSE9195** | Loi *et al.*, 2008 | 77 | 0 | | 0 | 0 |

| | | | | | | | |
|---|---|---|---|---|---|---|---|
| 1.0 | **GSE10510** | Calabrò *et al.*, 2009 | 139 | 134 | | 134 | 63 |
| 1.0 | **GSE11264** | Jézéquel *et al.*, 2009 | 252 | 252 | | 0 | 0 |
| 1.1 | **GSE11121** | Schmidt *et al.*, 2008 | 200 | 0 | | 0 | 0 |
| 1.1 | **GSE12093** | Zhang *et al.*, 2009 | 136 | 0 | | 0 | 0 |
| 3.1 | **GSE8757** | Chin *et al.*, 2007 | 171 | 171 | | 171 | 57 |
| 3.1 | **GSE7378** | Zhou *et al.*, 2007 | 54 | 0 | | 0 | 0 |
| 3.1 | **GSE16391** | Desmedt *et al.*, 2009 | 55 | 0 | | 0 | 0 |
| 3.1 | **GSE22133** | Jönsson *et al.*, 2010 | 346 | 339 | | 339 | 151 |
| 3.1 | **GSE19615** | Li *et al.*, 2010 | 115 | 0 | | 0 | 0 |
| 3.1 | **GSE17907** | Sircoulomb *et al.*, 2010 | 55 | 0 | | 0 | 0 |
| 3.1 | **GSE22219** | Buffa *et al.*, 2011 | 216 | 0 | | 0 | 0 |
| 3.1 | **GSE20711** | Dedeurwaerder *et al.*, 2011 | 85 | 0 | | 0 | 0 |
| 3.1 | **GSE26971** | Filipits *et al.*, 2011 | 277 | 0 | | 0 | 0 |
| 3.1 | **GSE25055** | Hatzis *et al.*, 2011 | 309 | 0 | | 0 | 0 |
| 3.1 | **GSE20685** | Kao *et al.*, 2011 | 296 | 296 | | 296 | 62 |
| 3.1 | **GSE21653** | Sabatier *et al.*, 2011 | 239 | 0 | | 0 | 0 |
| 3.1 | **GSE16987** | Wang *et al.*, 2011 | 149 | 0 | | 0 | 0 |
| 3.1 | **GSE45255** | Nagalla *et al.*, 2013 | 41 | 41 | | 41 | 10 |
| 4.3 | **GSE2109** | expO *et al.*, 2005 | 298 | 0 | | 0 | 0 |
| 4.3 | **GSE8193** | Yau *et al.*, 2007 | 47 | 0 | | 0 | 0 |
| 4.3 | **GSE20462** | Parris *et al.*, 2010 | 94 | 94 | | 94 | 44 |
| 4.3 | **GSE17705** | Symmans *et al.*, 2010 | 43 | 0 | | 0 | 0 |

| | | | | | | | |
|---|---|---|---|---|---|---|---|
| 4.3 | **GSE24450** | Heikkinen *et al.*, 2011 | 174 | 174 | | 174 | 27 |
| 4.3 | **GSE31448** | Sabatier *et al.*, 2011 | 71 | 0 | | 0 | 0 |
| 4.3 | **METABRIC** | Curtis *et al.*, 2012 | 1 980 | 1 980 | | 1 980 | 1 143 |
| 4.3 | **E_MTAB_365** | Guedj *et al.*, 2012 | 536 | 0 | | 0 | 0 |
| 4.3 | **GSE30682** | Servant *et al.*, 2012 | 343 | 0 | | 0 | 0 |
| 4.3 | **GSE42568** | Clarke *et al.*, 2013 | 104 | 104 | | 104 | 35 |
| 4.3 | **GSE40115** | Larsen *et al.*, 2013 | 183 | 0 | | 0 | 0 |
| 4.3 | **GSE55348** | Castagnoli *et al.*, 2014 | 53 | 0 | | 0 | 0 |
| 4.3 | **GSE43358** | Fumagalli *et al.*, 2014 | 56 | 55 | | 55 | 5 |
| 4.3 | **GSE36295** | Merdad *et al.*, 2014 | 45 | 0 | | 0 | 0 |
| 4.3 | **GSE37751** | Terunuma *et al.*, 2014 | 55 | 55 | | 55 | 19 |
| 4.3 | **GSE76274** | Burstein *et al.*, 2015 | 66 | 0 | | 0 | 0 |
| 4.3 | **GSE97177** | Biermann *et al.*, 2017 | 53 | 0 | | 0 | 0 |
| 4.5 | **GSE12276** | Bos *et al.*, 2009 | 204 | 204 | | 204 | 204 |
| 4.5 | **GSE18864** | Silver *et al.*, 2010 | 75 | 0 | | 0 | 0 |
| 4.6 | **GSE80999** | Aure *et al.*, 2017 | 381 | 0 | | 0 | 0 |
| 4.6 | **GSE86166** | Prabhakaran *et al.*, 2017 | 366 | 366 | | 366 | 103 |
| | | **Total:** | **10 872** | **5 622** | **52** | **5 113** | **2 231** |

**Summary:**

Targeted prognostic analyses for *CLPB* with all ER status, all PR status and all nodal status patients with overall survival (OS) information (n = 5 113).

**DNA microarray datasets which are used to analyze the disease-free survival rates by using bc-GenExMiner (v5.2)**

**Population and event data:**

| Ver | Study code | Original data | | Filtered data | | Final data | |
|---|---|---|---|---|---|---|---|
| | | Reference | No. patients | (ER all, PR all, N all and DFS) -selected patients | Gene found | No. patients | No. DFS |
| 1.0 | **Rosetta2002** | Van de Vijver *et al.*, 2002 | 295 | 295 | | 295 | 122 |
| 1.0 | **PNAS1732912100** | Sotiriou *et al.*, 2003 | 99 | 99 | | 0 | 0 |
| 1.0 | **GSE1379** | Ma *et al.*, 2004 | 59 | 59 | | 59 | 27 |
| 1.0 | **GSE2603** | Minn *et al.*, 2005 | 82 | 82 | | 82 | 27 |
| 1.0 | **GSE1456** | Pawitan *et al.*, 2005 | 159 | 159 | | 159 | 50 |
| 1.0 | **GSE2034** | Wang *et al.*, 2005 | 286 | 286 | | 286 | 107 |
| 1.0 | **GSE2741** | Weigelt *et al.*, 2005 | 50 | 50 | | 50 | 13 |
| 1.0 | **GSE3143** | Bild *et al.*, 2006 | 158 | 158 | | 0 | 0 |
| 1.0 | **E_TABM_158** | Chin *et al.*, 2006 | 112 | 112 | | 112 | 42 |
| 1.0 | **GSE4922** | Ivshina *et al.*, 2006 | 249 | 249 | | 249 | 89 |
| 1.0 | **GSE7390** | Desmedt *et al.*, 2007 | 198 | 198 | | 198 | 91 |
| 1.0 | **GSE6532** | Loi *et al.*, 2007 | 267 | 259 | | 259 | 88 |
| 1.0 | **GSE5327** | Minn *et al.*, 2007 | 58 | 58 | | 58 | 11 |
| 1.0 | **E_UCON_1** | Naderi *et al.*, 2007 | 135 | 135 | | 135 | 65 |
| 1.0 | **GSE7849** | Anders *et al.*, 2008 | 75 | 75 | | 0 | 0 |
| 1.0 | **GSE9893** | Chanrion *et al.*, 2008 | 151 | 151 | | 151 | 55 |
| 1.0 | **GSE9195** | Loi *et al.*, 2008 | 77 | 77 | | 77 | 13 |

| | | | | | | | |
|---|---|---|---|---|---|---|---|
| 1.0 | **GSE10510** | Calabrò *et al.*, 2009 | 139 | 134 | | 134 | 96 |
| 1.0 | **GSE11264** | Jézéquel *et al.*, 2009 | 252 | 252 | | 0 | 0 |
| 1.1 | **GSE11121** | Schmidt *et al.*, 2008 | 200 | 200 | | 200 | 46 |
| 1.1 | **GSE12093** | Zhang *et al.*, 2009 | 136 | 136 | | 136 | 20 |
| 3.1 | **GSE8757** | Chin *et al.*, 2007 | 171 | 171 | | 171 | 56 |
| 3.1 | **GSE7378** | Zhou *et al.*, 2007 | 54 | 54 | | 54 | 9 |
| 3.1 | **GSE16391** | Desmedt *et al.*, 2009 | 55 | 55 | | 55 | 55 |
| 3.1 | **GSE22133** | Jönsson *et al.*, 2010 | 346 | 339 | | 339 | 151 |
| 3.1 | **GSE19615** | Li *et al.*, 2010 | 115 | 115 | | 115 | 14 |
| 3.1 | **GSE17907** | Sircoulomb *et al.*, 2010 | 55 | 39 | | 39 | 17 |
| 3.1 | **GSE22219** | Buffa *et al.*, 2011 | 216 | 216 | | 216 | 82 |
| 3.1 | **GSE20711** | Dedeurwaerder *et al.*, 2011 | 85 | 85 | | 85 | 36 |
| 3.1 | **GSE26971** | Filipits *et al.*, 2011 | 277 | 258 | | 258 | 58 |
| 3.1 | **GSE25055** | Hatzis *et al.*, 2011 | 309 | 309 | | 309 | 65 |
| 3.1 | **GSE20685** | Kao *et al.*, 2011 | 296 | 296 | | 296 | 73 |
| 3.1 | **GSE21653** | Sabatier *et al.*, 2011 | 239 | 229 | | 229 | 74 |
| 3.1 | **GSE16987** | Wang *et al.*, 2011 | 149 | 147 | | 147 | 10 |
| 3.1 | **GSE45255** | Nagalla *et al.*, 2013 | 41 | 41 | | 41 | 14 |
| 4.3 | **GSE2109** | expO *et al.*, 2005 | 298 | 0 | | 0 | 0 |
| 4.3 | **GSE8193** | Yau *et al.*, 2007 | 47 | 0 | | 0 | 0 |
| 4.3 | **GSE20462** | Parris *et al.*, 2010 | 94 | 94 | | 94 | 44 |
| 4.3 | **GSE17705** | Symmans *et al.*, 2010 | 43 | 43 | | 43 | 8 |

| | | | | | | | |
|---|---|---|---|---|---|---|---|
| 4.3 | **GSE24450** | Heikkinen *et al.*, 2011 | 174 | 174 | | 174 | 34 |
| 4.3 | **GSE31448** | Sabatier *et al.*, 2011 | 71 | 0 | | 0 | 0 |
| 4.3 | **METABRIC** | Curtis *et al.*, 2012 | 1 980 | 1 980 | | 1 980 | 1 235 |
| 4.3 | **E_MTAB_365** | Guedj *et al.*, 2012 | 536 | 526 | | 526 | 118 |
| 4.3 | **GSE30682** | Servant *et al.*, 2012 | 343 | 0 | | 0 | 0 |
| 4.3 | **GSE42568** | Clarke *et al.*, 2013 | 104 | 104 | | 104 | 48 |
| 4.3 | **GSE40115** | Larsen *et al.*, 2013 | 183 | 0 | | 0 | 0 |
| 4.3 | **GSE55348** | Castagnoli *et al.*, 2014 | 53 | 53 | | 53 | 23 |
| 4.3 | **GSE43358** | Fumagalli *et al.*, 2014 | 56 | 55 | | 55 | 10 |
| 4.3 | **GSE36295** | Merdad *et al.*, 2014 | 45 | 0 | | 0 | 0 |
| 4.3 | **GSE37751** | Terunuma *et al.*, 2014 | 55 | 55 | | 55 | 19 |
| 4.3 | **GSE76274** | Burstein *et al.*, 2015 | 66 | 0 | | 0 | 0 |
| 4.3 | **GSE97177** | Biermann *et al.*, 2017 | 53 | 0 | | 0 | 0 |
| 4.5 | **GSE12276** | Bos *et al.*, 2009 | 204 | 204 | | 204 | 204 |
| 4.5 | **GSE18864** | Silver *et al.*, 2010 | 75 | 0 | | 0 | 0 |
| 4.6 | **GSE80999** | Aure *et al.*, 2017 | 381 | 0 | | 0 | 0 |
| 4.6 | **GSE86166** | Prabhakaran *et al.*, 2017 | 366 | 366 | | 366 | 119 |
| | | **Total:** | **10 872** | **9 232** | **52** | **8 648** | **3 538** |

### Summary:

Targeted prognostic analyses for *CLPB* with all ER status, all PR status and all nodal status patients with disease-free survival (DFS) information (metastatic or any relapse, or death) (n = 8 648).

**DNA microarray datasets which are used to analyze the distant metastasis-free survival rates by using bc-GenExMiner (v5.2)**

**Population and event data:**

| Ver | Study code | Original data | | Filtered data | | Final data | |
|---|---|---|---|---|---|---|---|
| | | Reference | No. patients | (ER all, PR all, N all and DMFS) -selected patients | Gene found | No. patients | No. DMFS |
| 1.0 | **Rosetta2002** | Van de Vijver *et al.*, 2002 | 295 | 295 | | 295 | 101 |
| 1.0 | **PNAS1732912100** | Sotiriou *et al.*, 2003 | 99 | 99 | | 0 | 0 |
| 1.0 | **GSE1379** | Ma *et al.*, 2004 | 59 | 0 | | 0 | 0 |
| 1.0 | **GSE2603** | Minn *et al.*, 2005 | 82 | 82 | | 82 | 27 |
| 1.0 | **GSE1456** | Pawitan *et al.*, 2005 | 159 | 159 | | 159 | 40 |
| 1.0 | **GSE2034** | Wang *et al.*, 2005 | 286 | 286 | | 286 | 107 |
| 1.0 | **GSE2741** | Weigelt *et al.*, 2005 | 50 | 50 | | 50 | 13 |
| 1.0 | **GSE3143** | Bild *et al.*, 2006 | 158 | 0 | | 0 | 0 |
| 1.0 | **E_TABM_158** | Chin *et al.*, 2006 | 112 | 112 | | 112 | 21 |
| 1.0 | **GSE4922** | Ivshina *et al.*, 2006 | 249 | 0 | | 0 | 0 |
| 1.0 | **GSE7390** | Desmedt *et al.*, 2007 | 198 | 198 | | 198 | 62 |
| 1.0 | **GSE6532** | Loi *et al.*, 2007 | 267 | 259 | | 259 | 66 |
| 1.0 | **GSE5327** | Minn *et al.*, 2007 | 58 | 58 | | 58 | 11 |
| 1.0 | **E_UCON_1** | Naderi *et al.*, 2007 | 135 | 0 | | 0 | 0 |
| 1.0 | **GSE7849** | Anders *et al.*, 2008 | 75 | 75 | | 0 | 0 |
| 1.0 | **GSE9893** | Chanrion *et al.*, 2008 | 151 | 151 | | 151 | 46 |
| 1.0 | **GSE9195** | Loi *et al.*, 2008 | 77 | 77 | | 77 | 10 |

| | | | | | | | |
|---|---|---|---|---|---|---|---|
| 1.0 | **GSE10510** | Calabrò *et al.*, 2009 | 139 | 0 | | 0 | 0 |
| 1.0 | **GSE11264** | Jézéquel *et al.*, 2009 | 252 | 252 | | 0 | 0 |
| 1.1 | **GSE11121** | Schmidt *et al.*, 2008 | 200 | 200 | | 200 | 46 |
| 1.1 | **GSE12093** | Zhang *et al.*, 2009 | 136 | 136 | | 136 | 20 |
| 3.1 | **GSE8757** | Chin *et al.*, 2007 | 171 | 171 | | 171 | 38 |
| 3.1 | **GSE7378** | Zhou *et al.*, 2007 | 54 | 54 | | 54 | 9 |
| 3.1 | **GSE16391** | Desmedt *et al.*, 2009 | 55 | 0 | | 0 | 0 |
| 3.1 | **GSE22133** | Jönsson *et al.*, 2010 | 346 | 0 | | 0 | 0 |
| 3.1 | **GSE19615** | Li *et al.*, 2010 | 115 | 115 | | 115 | 14 |
| 3.1 | **GSE17907** | Sircoulomb *et al.*, 2010 | 55 | 39 | | 39 | 17 |
| 3.1 | **GSE22219** | Buffa *et al.*, 2011 | 216 | 216 | | 216 | 82 |
| 3.1 | **GSE20711** | Dedeurwaerder *et al.*, 2011 | 85 | 0 | | 0 | 0 |
| 3.1 | **GSE26971** | Filipits *et al.*, 2011 | 277 | 258 | | 258 | 58 |
| 3.1 | **GSE25055** | Hatzis *et al.*, 2011 | 309 | 309 | | 309 | 65 |
| 3.1 | **GSE20685** | Kao *et al.*, 2011 | 296 | 296 | | 296 | 63 |
| 3.1 | **GSE21653** | Sabatier *et al.*, 2011 | 239 | 0 | | 0 | 0 |
| 3.1 | **GSE16987** | Wang *et al.*, 2011 | 149 | 0 | | 0 | 0 |
| 3.1 | **GSE45255** | Nagalla *et al.*, 2013 | 41 | 41 | | 41 | 14 |
| 4.3 | **GSE2109** | expO *et al.*, 2005 | 298 | 0 | | 0 | 0 |
| 4.3 | **GSE8193** | Yau *et al.*, 2007 | 47 | 0 | | 0 | 0 |
| 4.3 | **GSE20462** | Parris *et al.*, 2010 | 94 | 0 | | 0 | 0 |
| 4.3 | **GSE17705** | Symmans *et al.*, 2010 | 43 | 43 | | 43 | 8 |

| | | | | | | | |
|---|---|---|---|---|---|---|---|
| 4.3 | **GSE24450** | Heikkinen *et al.*, 2011 | 174 | 174 | | 174 | 34 |
| 4.3 | **GSE31448** | Sabatier *et al.*, 2011 | 71 | 0 | | 0 | 0 |
| 4.3 | **METABRIC** | Curtis *et al.*, 2012 | 1 980 | 1 980 | | 1 980 | 602 |
| 4.3 | **E_MTAB_365** | Guedj *et al.*, 2012 | 536 | 526 | | 526 | 118 |
| 4.3 | **GSE30682** | Servant *et al.*, 2012 | 343 | 0 | | 0 | 0 |
| 4.3 | **GSE42568** | Clarke *et al.*, 2013 | 104 | 0 | | 0 | 0 |
| 4.3 | **GSE40115** | Larsen *et al.*, 2013 | 183 | 0 | | 0 | 0 |
| 4.3 | **GSE55348** | Castagnoli *et al.*, 2014 | 53 | 0 | | 0 | 0 |
| 4.3 | **GSE43358** | Fumagalli *et al.*, 2014 | 56 | 55 | | 55 | 10 |
| 4.3 | **GSE36295** | Merdad *et al.*, 2014 | 45 | 0 | | 0 | 0 |
| 4.3 | **GSE37751** | Terunuma *et al.*, 2014 | 55 | 0 | | 0 | 0 |
| 4.3 | **GSE76274** | Burstein *et al.*, 2015 | 66 | 0 | | 0 | 0 |
| 4.3 | **GSE97177** | Biermann *et al.*, 2017 | 53 | 0 | | 0 | 0 |
| 4.5 | **GSE12276** | Bos *et al.*, 2009 | 204 | 0 | | 0 | 0 |
| 4.5 | **GSE18864** | Silver *et al.*, 2010 | 75 | 0 | | 0 | 0 |
| 4.6 | **GSE80999** | Aure *et al.*, 2017 | 381 | 0 | | 0 | 0 |
| 4.6 | **GSE86166** | Prabhakaran *et al.*, 2017 | 366 | 0 | | 0 | 0 |
| | | **Total:** | **10 872** | **6 766** | **52** | **6 340** | **1 702** |

**Summary:**

Targeted prognostic analyses for *CLPB* with all ER status, all PR status and all nodal status patients with distant metastasis-free survival (DMFS) information (n = 6 340).

**RNA-Seq datasets which are used to analyze the overall survival rates by using bc-GenExMiner (v5.2)**

**Population and event data:**

| Ver | Study code | Original data | | Filtered data | | Final data | |
|---|---|---|---|---|---|---|---|
| | | Reference | No. patients | (ER all, PR all, N all and OS) -selected patients | Gene found | No. patients | No. OS |
| 4.3 | **TCGA** | TCGA *et al.*, 2012 | 743 | 743 | | 743 | 95 |
| 4.3 | **SCAN-B / GSE96058** | Saal *et al.*, 2015 | 3 273 | 3 273 | | 3 273 | 336 |
| 4.3 | **SCAN-B / GSE81538** | Brueffer *et al.*, 2018 | 405 | 0 | | 0 | 0 |
| | | **Total:** | **4 421** | **4 016** | **3** | **4 016** | **431** |

**Summary:**

Targeted prognostic analyses for *CLPB* with all ER status, all PR status and all nodal status patients with overall survival (OS) information (n = 4 016).

**RNA-Seq datasets which are used to analyze the disease-free survival rates by using bc-GenExMiner (v5.2)**

**Population and event data:**

| Ver | Study code | Original data | | Filtered data | | Final data | |
|---|---|---|---|---|---|---|---|
| | | Reference | No. patients | (ER all, PR all, N all and DFS) -selected patients | Gene found | No. patients | No. DFS |
| 4.3 | **TCGA** | TCGA *et al.*, 2012 | 743 | 743 | | 743 | 131 |
| 4.3 | **SCAN-B / GSE96058** | Saal *et al.*, 2015 | 3 273 | 3 273 | | 3 273 | 336 |
| 4.3 | **SCAN-B / GSE81538** | Brueffer *et al.*, 2018 | 405 | 0 | | 0 | 0 |
| | | **Total:** | **4 421** | **4 016** | **3** | **4 016** | **467** |

**Summary:**

Targeted prognostic analyses for *CLPB* with all ER status, all PR status and all nodal status patients with disease-free survival (DFS) information (metastatic or any relapse, or death) (n = 4 016).

**RNA-Seq datasets which are used to analyze the distant metastasis-free survival rates by using bc-GenExMiner (v5.2)**

**Population and event data:**

| Ver | Study code | Original data | | Filtered data | | Final data | |
|---|---|---|---|---|---|---|---|
| | | Reference | No. patients | (ER all, PR all, N all and DMFS) -selected patients | Gene found | No. patients | No. DMFS |
| 4.4 | **TCGA** | TCGA *et al.*, 2012 | 743 | 743 | | 743 | 41 |
| | | **Total:** | **743** | **743** | **1** | **743** | **41** |

**Summary:**

Targeted prognostic analyses for *CLPB* with all ER status, all PR status and all nodal status patients with distant metastasis-free survival (DMFS) information (n = 743).
